# ArlCDE form the archaeal switch complex

**DOI:** 10.1101/2020.04.09.033365

**Authors:** Zhengqun Li, Marta Rodriguez-Franco, Sonja-Verena Albers, Tessa E. F. Quax

## Abstract

Cells require a sensory system and a motility structure to achieve directed movement. Bacteria and archaea both possess rotating filamentous motility structures that work in concert with the sensory chemotaxis system. This allows microorganisms to move along chemical gradients. The central response regulator protein CheY can bind to the motor of the motility structure, the flagellum in bacteria and the archaellum in archaea. Both motility structures have a fundamentally different protein composition and structural organization. Yet, both systems receive input from the chemotaxis system. We applied a fluorescent microscopy approach in the model euryarchaeon *Haloferax volcanii*, and shed light on the sequence order in which signals are transferred from the chemotaxis system to the archaellum. Our findings indicate that the euryarchaeal specific ArlCDE are part of the archaellum motor and that they directly receive input from the chemotaxis system via the adaptor protein CheF. Hence, ArlCDE are an important feature of the archaellum of euryarchaea, are essential for signal transduction during chemotaxis and represent the archaeal switch complex.

## Introduction

Bacteria and Archaea employ rotating filamentous structures to swim through liquid environments. Although they are functionally similar, there is no structural similarity between the filaments from Bacteria and Archaea (Berg and Anderson, 1973; Alam and Oesterhelt, 1984; Chevance and Hughes, 2008; Jarrell and Albers, 2012; Albers and Jarrell, 2018; Beeby *et al.*, 2020). The archaeal motility structure, the archaellum, has homology to bacterial type IV pili, and it consist of ∼10 Arl proteins (previously named Fla proteins) (Kalmokoff and Jarrell, 1991; Jarrell and Albers, 2012; Pohlschroder *et al.*, 2018). The energy required for rotation is derived from ATP hydrolysis (Thomas *et al.*, 2002; Streif *et al.*, 2008; Reindl *et al.*, 2013), and new protein subunits are N-terminally processed and added at the base of the growing filament (Kalmokoff and Jarrell, 1991; Bardy and Jarrell, 2003; Szabo *et al.*, 2006). In contrast, the bacterial motility structure, the flagellum, consists of over 30 different proteins, which are not found in archaea (Chevance and Hughes, 2008; Erhardt *et al.*, 2010; Lassak *et al.*, 2012; Altegoer and Bange, 2015). The energy for flagellum rotation originates from the proton motive force and new protein subunits are added at the tip of the growing filament (Chevance and Hughes, 2008; Erhardt *et al.*, 2010; Altegoer and Bange, 2015).

Despite the fundamental differences in the organization of the archaeal and bacterial motility structures, Archaea and Bacteria both possess the chemotaxis system (Briegel *et al.*, 2015; Quax *et al.*, 2018a). This system allows cells to move along chemical gradients and, in combination with the motility structure, ensures directional movement (Szurmant and Ordal, 2004; Porter *et al.*, 2011; Bi and Sourjik, 2018). Archaea, such as euryarchaea and some thaumarchaea, have likely received the chemotaxis system from bacteria via horizontal gene transfer (Wuichet *et al.*, 2010; Wuichet and Zhulin, 2010; Briegel *et al.*, 2015).

In bacteria the chemotaxis system transfers signals to the base of the flagellum (Sourjik and Berg, 2002; Chevance and Hughes, 2008). Attractants or repellents can bind to chemosensory receptors, methyl-accepting chemotaxis proteins (MCPs), which are often presented at the cell surface (Parkinson *et al.*, 2015; Salah Ud-Din and Roujeinikova, 2017). These MCPs are organized together with CheW and CheA proteins in chemosensory arrays, which are large organized clusters that ensure signal integration and amplification (Briegel *et al.*, 2012; Briegel *et al.*, 2014). Binding of stimuli to the MCPs triggers a signaling cascade, which eventually leads to phosphorylation of the response regulator protein CheY (Welch *et al.*, 1993; Parkinson *et al.*, 2015). Phosphorylated CheY binds with higher affinity to the switch complex at the flagellum motor, which consists of FliM, FliN, and FliG proteins (Barak and Eisenbach, 1992; Sarkar *et al.*, 2010; Paul *et al.*, 2011). This results in a change or a pause in flagellum rotation direction (Sarkar *et al.*, 2010; Paul *et al.*, 2011). These bacterial switch proteins are, like the other flagellum proteins, absent from archaea (Jarrell and Albers, 2012) and the archaeal equivalent of a switch complex has not yet been identified. Therefore, it is still an open question how the archaeal motility structure receives signals from the bacterial-like chemotaxis system.

Recently available cryo-EM structures of the archaellum provide clues to this question. Sub-tomogram averaging of the archaellum motor of the euryarchaeon *Pyrococcus furiosus*, revealed a bell-like structure below the motor that stretches into the cytoplasm (Daum *et al.*, 2017). The central core of the motor is formed by the proteins ArlJ, ArlI and ArlH (previously named FlaJ, I and H) encoded from the archaellum operon (Figure 1a) (Ghosh *et al.*, 2011; Banerjee *et al.*, 2013; Reindl *et al.*, 2013; Chaudhury *et al.*, 2016; Albers and Jarrell, 2018; Chaudhury *et al.*, 2018). ArlJ is an integral membrane protein that was not well resolved in the cryo-Em structure (Daum *et al.*, 2014). Six copies of the cytosolic ATPase ArlI and six of the ATP binding protein ArlH, could be mapped into the cytosolic part of the central core of the archaellum (Daum *et al.*, 2017). After mapping I and H, still a large proportion of the cytosolic part of the motor has remained unassigned (Daum *et al.*, 2017). This density has a ring-like shape and was hypothesized to be occupied by ArlC, D and E, because these are the only proteins with unassigned function encoded in eurayarchaeal archaellum operons (Figure 1a). It was suggested that these proteins together form a ring structure, in analogy with the ring structure formed by the crenarchaeal specific protein FlaX protein (Daum *et al.*, 2017; Briegel *et al.*, 2017). In many archaeal genomes, different combinations of fusions of ArlC, D and E are encoded, such as ArlCE from the halophilic euryarchaeon *Haloferax volcanii*, which suggests that the three proteins might form a complex (Albers and Jarrell, 2015). Since the unassigned ring-like density is at the most peripheral part of the motor, it would represent a convenient docking place for chemotaxis proteins. Corresponding with this hypothesis, an interactome study in *Halobacterium salinarum*, previously indicated that ArlCE (fused in this organism) interacts with the archaeal specific chemotaxis protein CheF (Figure 1a) (Schlesner *et al.*, 2009; Schlesner *et al.*, 2012). Motile archaea encoding the bacterial-like chemotaxis system, all possess the adaptor protein CheF, which can bind to CheY and as such forms a link between the chemotaxis system and the archaellum (Schlesner *et al.*, 2009; Quax *et al.*, 2018b).

**Figure 1.**
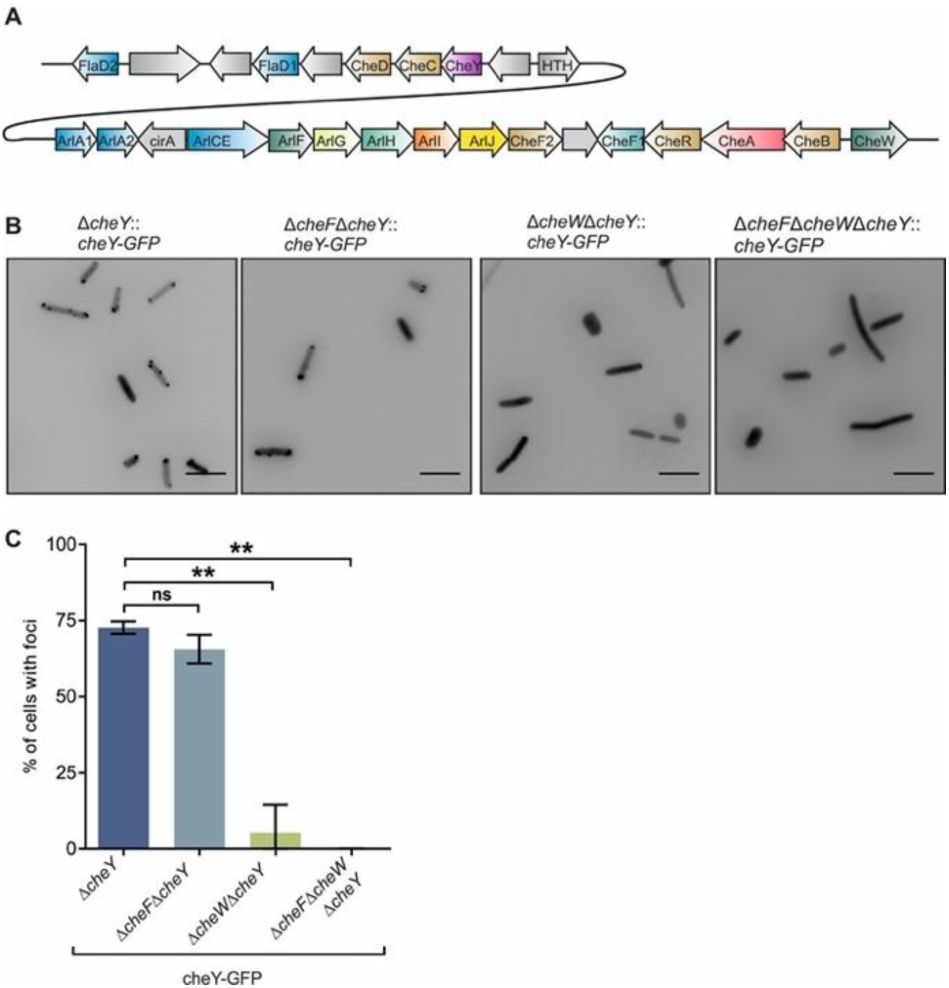
Intracellular positioning of CheY is mainly dependent on chemosensory arrays in *H. volcanii*. (a) Schematic representation of the archaellum and chemotaxis operon in the model euryarchaeal *H. volcanii* (b) Representative fluorescent micrographs of the intracellular distribution of CheY-GFP clusters in different *H. volcanii* mutants in early stationary phase. I, Δ*cheY*; II, Δ*cheF* Δ*cheY*; III, Δ*cheW* Δ*cheY*; IV, Δ*cheF* Δ*cheW* Δ*cheY*. The number of cells analyzed for each mutant is n>500. Scale bars, 5 µm. (c) Percentages of cells with CheY-GFP foci in the four strains analyzed in A. ** *P* < 0.01. ns, not significant *P* > 0.05.

We aimed to establish in which sequence order chemotaxis proteins transfer signals to the archaellum motor and which motor proteins are likely in direct contact with the chemotaxis system. To address these questions, we used *Haloferax volcanii*, for which a well-developed genetic manipulation system is available, in addition to previously obtained information on the cellular positioning of several chemotaxis and archaellum proteins (Allers *et al.*, 2004; Li *et al.*, 2019). Fluorescent microscopy was employed to study the localization of several proteins, important for signal transduction between the chemotaxis system and the motility machinery. This work sheds light onto the role of the ArlC, D, and E archaellum proteins and indicates that they are a crucial factor in receiving input from the chemotaxis system.

## Results

### The chemosensory arrays anchor the archaeal response regulator CheY

We aimed to gain insight into the sequence in which archaeal chemotaxis and archaellum proteins are interacting with each other. First, we focused on the response regulator CheY, which is thought to shuttle between the chemotaxis system and the motility structure (Figure 1a). Previously, we have shown that CheY localizes primarily to the cell poles of motile cells, but it is also present at the lateral membranes (Li *et al.*, 2019). This positioning pattern shares similarities with that of the chemosensory arrays and with that of archaella, which are exclusively present at cell poles (Li *et al.*, 2019). As CheY was shown to interact with the archaeal specific chemotaxis protein CheF (Schlesner *et al.*, 2009; Quax *et al.*, 2018b), we wondered if it might position at the archaellum motor in a CheF dependent manner. A *H. volcanii* Δ*cheY* Δ*cheF* mutant was constructed in which either CheY-GFP or GFP-CheF were expressed. These fusion proteins were shown to be correctly expressed by Western blot analysis and were previously shown to restore motility on semi-solid agar plates (Figure S1a. Table S1) (Li *et al.*, 2019). The positioning pattern of GFP-CheY did not significantly differ in the Δ*cheY*Δ*cheF* and in the *ΔcheY* strain (Figure 1b,c). In both cases, polar and lateral foci were observed, suggesting that localization of the response regulator CheY is mainly independent of CheF (Figure 1b, c).

Next, we tested if CheY might bind to chemosensory arrays. CheW was previously used as a marker protein to indicate the cellular position of chemosensory arrays in *H. volcanii* (Li *et al.*, 2019). A Δ*cheY* Δ*cheW* deletion strain was created in which CheY-GFP was expressed. Observation of this strain by fluorescent microscopy showed that cells mostly displayed a diffuse fluorescent signal in the cytoplasm, with the exception of a few polar foci in a low number of cells (5.3%) (Figure 1b, c). When GFP-CheY was expressed in a Δ*cheY*Δ*cheF*Δ*cheW* strain, also the residual signal at the cell poles disappeared and the fluorescent signal was exclusively diffuse (Figure 1b, c). These findings indicate that in exponentially growing cells, CheY is mainly bound to chemosensory arrays, while a very small fraction might be present at the archaellum motor via binding to CheF.

### ArlD and ArlCE are important for communication with the chemotaxis system

CheF was previously shown to position mainly at the cell poles of *H. volcanii* (Li *et al.*, 2019). Therefore, we asked if CheF could be permanently localized at the archaellum motor in motile cells. The proteins ArlD and ArlCE both are encoded in the archaellum operon in euryarchaea and have an unknown function (Figure 1a) (Albers and Jarrell, 2015; Albers and Jarrell, 2018). They are hypothesized to be part of the archaellum motor (Daum *et al.*, 2017; Briegel *et al.*, 2017) and preliminary data from *M. maripaludis* suggest that they are membrane associated, although they both lack a transmembrane domain (Thomas and Jarrell, 2001; Thomas *et al.*, 2002). As an interactome study suggested that ArlCE might be in direct contact with the archaeal specific chemotaxis protein CheF (Schlesner *et al.*, 2009), we asked if CheF requires ArlCE and ArlD to bind to the archaellum motor. As *H. volcanii* encodes two ArlD homologs, we made knock-outs of both genes. *arlD2* is encoded from a gene a little upstream of the archaellum operon (Figure 1a). Analysis of a Δ*arlD2* strain on semi-solid agar plate, showed that directional movement and motility were comparable to the wild type (Figure S2a, b). In contrast, Δ*arlD1* has a severe motility defect, as the archaellum is not correctly produced anymore (Li *et al.*, 2019). Therefore, we continued with ArlD1 and refer to this protein as ArlD, throughout the paper. Next, we constructed the Δ*arlD* Δ*cheF* and Δ*arlCE* Δ*cheF* deletion strains in *H. volcanii*. GFP-CheF was expressed in Δ*arlD* Δ*cheF* and Δ*arlCE* Δ*cheF*. Interestingly, the positioning pattern of GFP-CheF showed significant difference in both a Δ*arlD* Δ*cheF* and a Δ*arlCE* Δ*cheF* strain compared to that in a Δ*cheF* strain (Figure 2a, b). GFP-CheF foci were observed at the cell poles in 65% of Δ*cheF* cells, consistent with what has been described previously (Table S1) (Li *et al.*, 2019). In contrast, the number of cells with polar GFP-CheF foci was significantly reduced in both the Δ*arlD* Δ*cheF* and the Δ*arlCE* Δ*cheF* strain (to ∼20% of cells) (Figure 2a, b). These findings indicate that ArlD and ArlCE promote, but are not essential for, CheF positioning to the cell pole. It might be possible that the largest fraction of CheF proteins is positioned at the cell pole via interaction with ArlCE and ArlD that are bound to the archaellum motor. A smaller fraction of CheF might be positioned at the cell pole because of other protein interactions.

**Figure 2.**
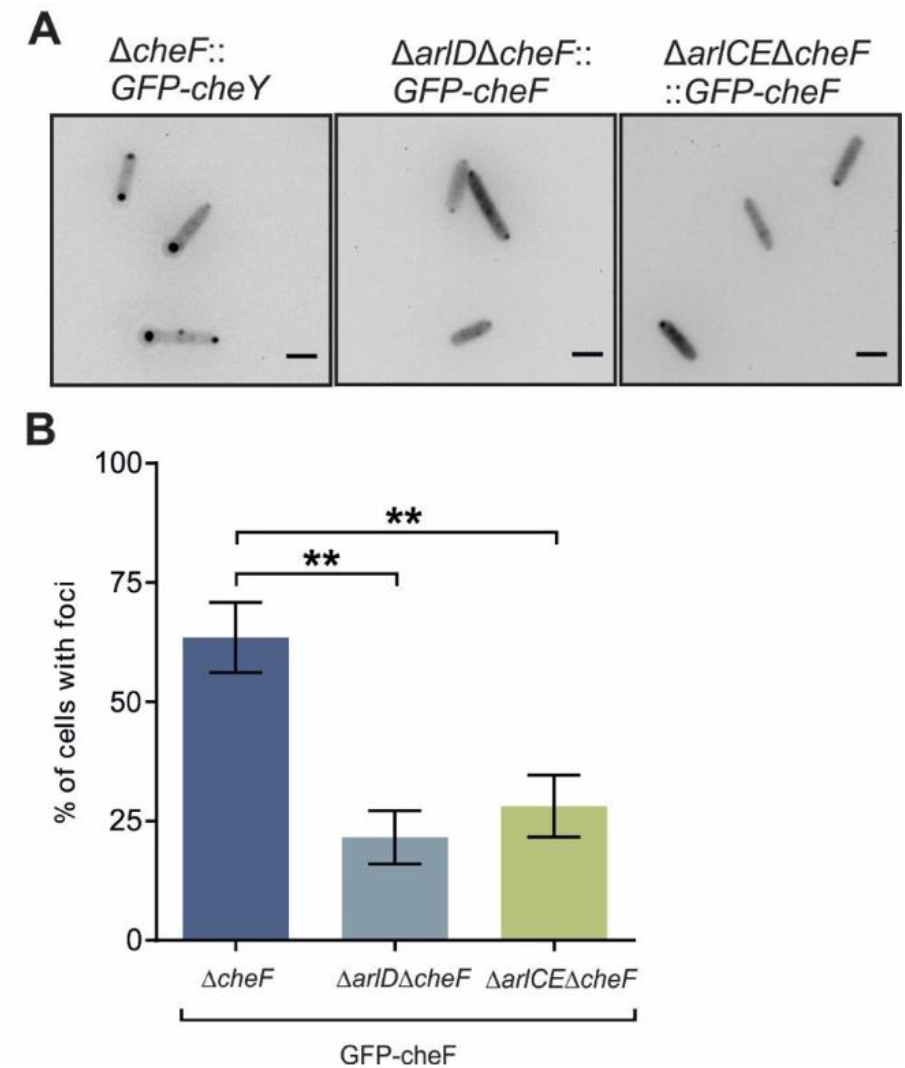
The archaellum proteins ArlD and ArlCE are important for communication with the chemotaxis system. (a) Representative fluorescent images of the intracellular distribution of GFP-CheF clusters in different *H. volcanii* mutants in exponential phase Scale bars, 2 µm. (b) Percentages of cells with GFP-CheF clusters in the strains described in (a). The number of cells of each mutant analyzed is n > 500. \*\**P* < 0.01 as established by T-test.

### Polar positioning of ArlD and ArlCE is inter-dependent

After we demonstrated the importance of ArlD and ArlCE for positioning of the chemotaxis adaptor protein CheF, we aimed to gain more information on the possible function of these two proteins. Previously we have shown that ArlD is likely part of the archaellum motor, as fluorescent fusion proteins of ArlD were positioned at the cell poles of rod-shaped motile *H. volcanii* cells, the location where also archaella are found (Table S1) (Li *et al.*, 2019). In order to test if the ArlCE protein is also part of the archaellum motor, we constructed a Δ*arlCE* deletion strain. Analysis by EM showed that this strain does not have archaella at its surface (Figure 3a), and did not form motility rings on semi-solid agar plate (Figure 3b). We expressed N- and C-terminal GFP fusions of ArlCE in a Δ*arlCE* strain. While expression of native ArlCE could restore the motile phenotype on semi-solid agar plate (Figure 3b), both GFP fusions did not restore motility (Table S1). Western blot analysis with α-GFP antibodies did not show a clear signal for either fusion, indicating that expression of ArlCE fusions to GFP yielded only low levels of the fusion proteins (data not shown). Correspondingly, Δ*arlCE* strains expressing GFP-ArlCE or ArlCE-GFP observed by fluorescence microscopy, had only a very low total GFP signal. Still, distinct foci were observed, exclusively near the cell poles of the motile rod-shaped cells (Figure 3c). 39% of the cells showed an ArlCE-GFP signal at one pole, while 42% showed bipolar ArlCE-GFP foci. The percentages were similar for the GFP-ArlCE signals. This positioning pattern is reminiscent of that of ArlD, suggesting that ArlCE also positions at the archaellum motor (Li *et al.*, 2019).

**Figure 3.**
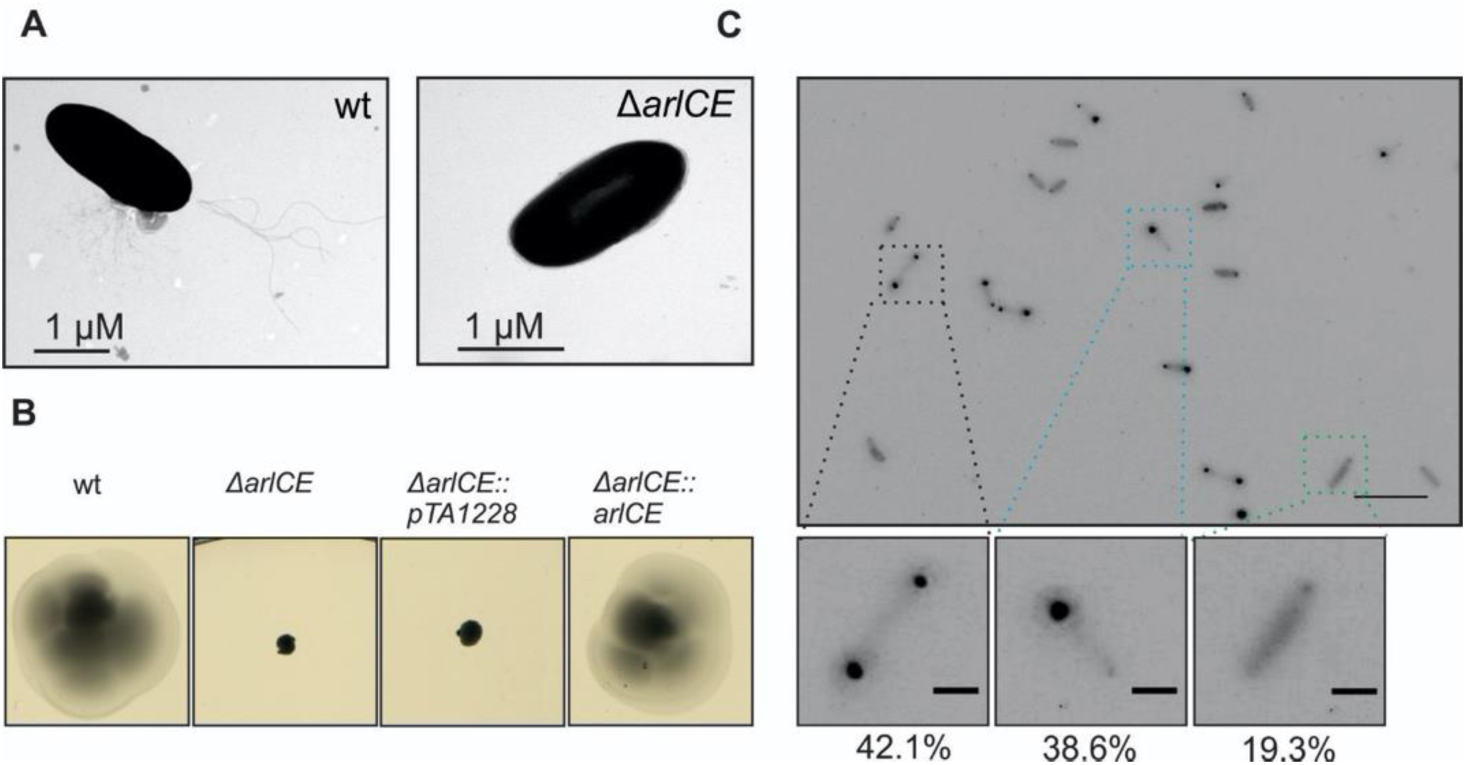
The positioning pattern of ArlCE is reminiscent of that of ArlD. (a) Transmission electron microscopy (TEM) of the wild-type (WT) and Δ*arlCE H. volcanii* cells. Scale bar, 1 µm. (b) Influence of ArlCE on directional movement. Representative image of motility assay of different *H. volcanii* strains on semis solid agar plates. This experiment was performed on at least 3 independent occasions. pTA1228, empty plasmid (c) Representative fluorescent image of ArlCE-GFP clusters in *H.volcanii* Δ*arlCE* cells in early exponential phase. The percentages of cells with each positioning pattern are showed below the corresponding exemplary images. The total population of the analyzed cells was n > 500. Scale bars, 10 µm (upper panel) and 2 µm (lower panels).

To confirm this hypothesis, ArlD-GFP and ArlCE-mCherry were co-expressed in a Δ*arlD* Δ*arlCE* strain. The signals of both proteins overlapped, suggesting ArlD and ArlCE co-localized (Figure 4a). To study if the positioning of the two proteins was interdependent, ArlD-GFP or ArlCE-GFP were expressed in the Δ*arlCE* Δ*arlD* strain. Western blot analysis showed that ArlD-GFP was correctly expressed in this background (Figure S1c). Comparison with the positioning pattern of the two proteins in the control strains, showed that the number of cells with distinct ArlD or ArlCE foci, was significantly reduced in absence of either of the other protein (Figure 4b, S2c). Thus, the correct positioning of ArlD or ArlCE at the archaellum motor is dependent on the presence of both proteins. This suggests that ArlD and ArlCE might form a precomplex, required for binding to the archaellum motor.

**Figure 4.**
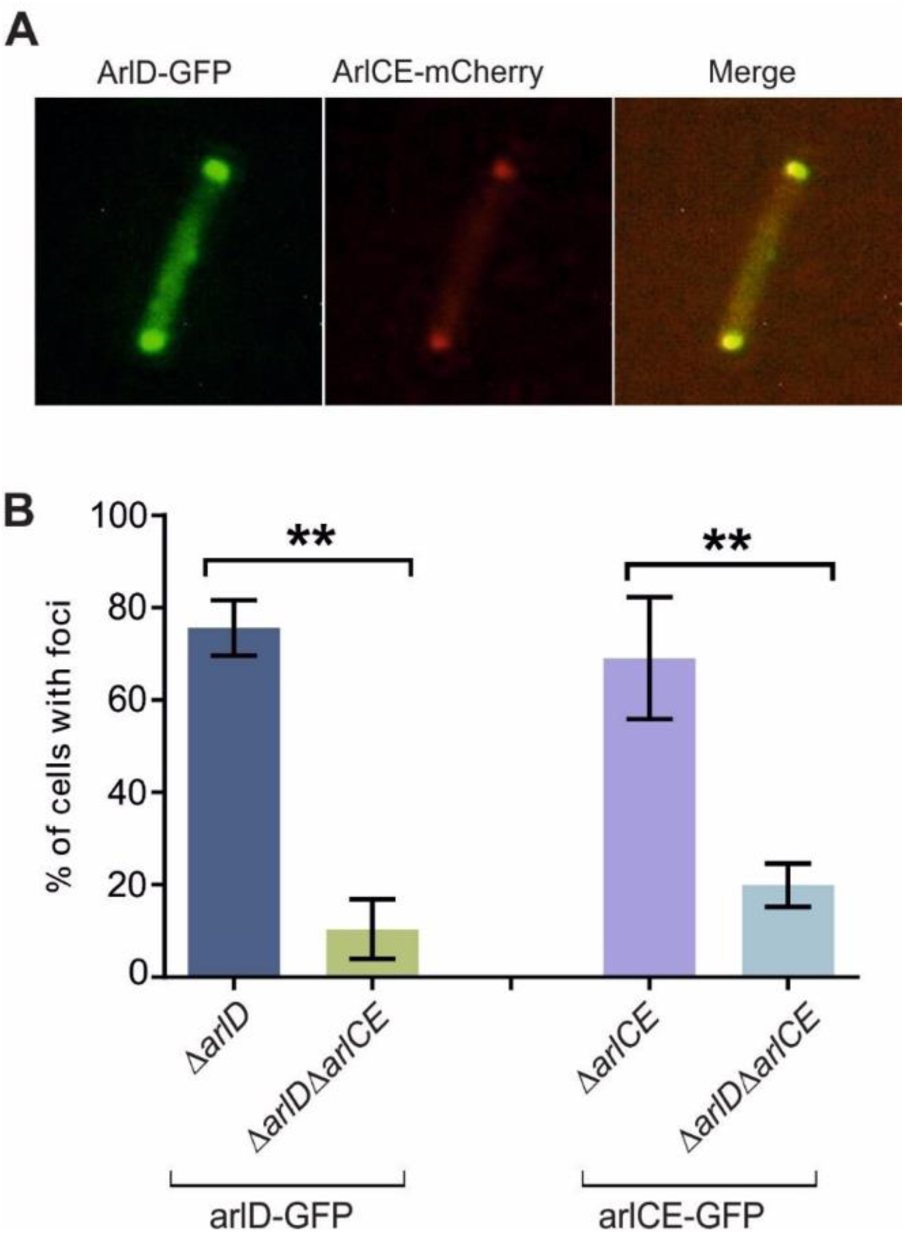
The positioning of ArlD and ArlCE are dependent on each other in *H.volcanii*. (a) Co-expression of ArlD-GFP and ArlCE-mCherry in the *H.volcanii* Δ*arlD* Δ*arlCE* strain in early exponential phase. (b) Percentages of cells with intracellular ArlD or ArlCE foci in single and double knockout strains. N>500. \*\**P* < 0.01 as established by T-test.

ArlD and ArlCE promote positioning of CheF at the cell pole (Figure 2). To test if ArlD and ArlCE also require CheF for polar localization, we expressed ArlD-GFP and ArlCE-GFP in a Δ*arlD* Δ*cheF* strain and in a Δ*arlCE* Δ*cheF* strain, respectively. We compared the number of cells with polar foci in these strains with those of the Δ*arlD* ArlD-GFP and the Δ*arlCE* ArlCE-GFP strain. The positioning pattern of ArlD and ArlCE was not significantly affected in the absence CheF (Figure S3). Thus, ArlD and ArlCE do not require CheF for correct positioning at the cell pole.

### ArlD and ArlCE dock on the archaellum motor via binding to ArlH

As we demonstrated that CheF is not responsible for polar positioning of ArlCE and ArlD, we hypothesized that the ArlCE and ArlD proteins are likely localized at the cell pole by direct binding to the archaellum motor. The core of the motor consists of ArlH, ArlI and the membrane protein ArlJ, which interact with each other to form a large oligomeric complex (Banerjee *et al.*, 2013; Reindl *et al.*, 2013; Chaudhury *et al.*, 2016). The cryo-EM structure of the archaellum motor of euryarchaea was resolved previously, which indicated that ArlH interacts with a ring like density that might consist of ArlC, D and E (Daum *et al.*, 2017). Therefore, we first tested if ArlD and ArlCE might require ArlH for polar localization. First, a Δarl*H* strain was constructed in *H. volcanii*. Analysis on semi-solid agar plate showed that the Δarl*H* strain did not form motility rings, in correspondence with the fact that a mutant in other archaea, was previously shown to possess no archaella (Chaban *et al.*, 2007; Staudinger, 2008; Chaudhury *et al.*, 2016) (Figure 5a). Expression of native ArlH in the Δ*arlH* strain could complement the ability of motility on semi-solid agar plate (Figure 5a). In contrast, expression of an ArlH-GFP fusion did not restore motility (Table S1). However, Western blot analysis indicated that the fusion protein ArlH-GFP was correctly expressed (Figure S1b) and expression of ArlH-GFP in the Δ*arlH* strain resulted in foci at the cell pole in the majority of cells (64%), corresponding with fact that ArlH is part of the archaellum motor (Figure 5b). Possibly, fusion of GFP to the C-terminus of ArlH, blocks interaction with other proteins or hinders its function, thus rendering this mutant non-motile.

**Figure 5.**
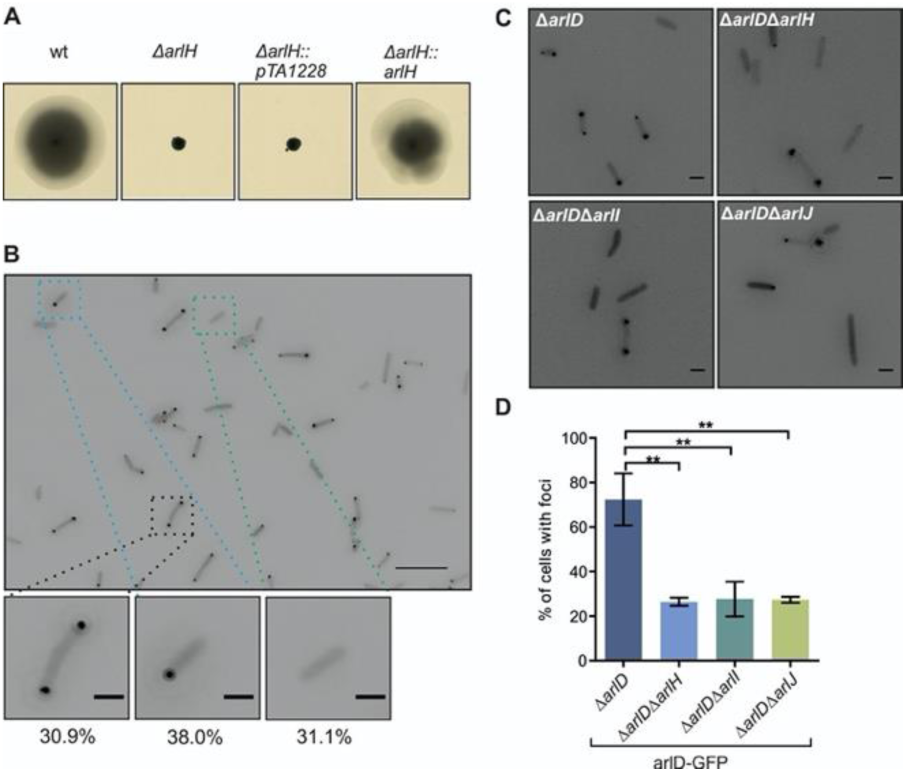
Polar localization of ArlD is mainly determined by binding to the archaellum motor. (a) Influence of ArlH on directional movement. Motility assay on semi-solid agar plate of different *H. volcanii* strains in rich medium. pTA1228, empty plasmid (b) Intracellular distribution of GFP-ArlH clusters in the *H. volcanii* Δ*arlH* strain. The percentage of total cells with each pattern are showed at the bottom. n > 500. Scale bars, 10 µm (upper panel) and 2 µm (lower panels). (c) Representative fluorescent images of intracellular distribution of ArlD-GFP *H. volcanii* strains in which different archaellum motor proteins were deleted. Δ*arlD*, Δ*arlD* Δ*arlH*, Δ*arlD* Δ*arlI*, Δ*arlD* Δ*arlJ*. The number of each analyzed mutant, n>500. Scale bars, 2 µm. (d) Percentages of cells with intracellular ArlD-GFP foci in strains described in (c). ** p< 0.01 as established by T-test.

Western blot analysis of the GFP-arlH fusion protein showed two bands, in size corresponding to ArlH and to GFP (Figure S1b). Thus, the GFP-ArlH fusion protein was likely cleaved. Fluorescence microscopy analysis suggested that this is indeed the case, as GFP-ArlH expression in a Δ*arlH* resulted in a diffuse GFP signal in the cytoplasm (Figure 5b). GFP-ArlH could restore the motility of a Δ*arlH* strain on semi-solid agar plate, which is likely the result of incorporation of the cleaved ArlH, without GFP tag (Figure 5a), in the archaellum motor.

Next, the double deletion strains Δ*arlD* Δ*arlH* and Δ*arlCE* Δ*arlH* were constructed to test the influence of ArlH on ArlD and ArlCE positioning. The number of cells with polar foci was reduced significantly in the Δ*arlD* Δ*arlH* strain expressing ArlD-GFP (27%), in comparison with expression of this construct in the Δ*arlD* strain (69%)(Figure 5c, d). ArlD-GFP was correctly expressed in this background as observed by Western blot analysis (Figure S1c). A similar observation was made for ArlCE-GFP expression in the Δ*arlCE* Δ*arlH* strain, where a reduction of ∼70% to ∼36% of cells with polar foci was detected (Figure S4a, b). As the polar positioning of both ArlD and ArlCE was severely diminished in absence of ArlH (Figure 5c, d, S4a, b), ArlD and ArlCE likely dock to the archaellum motor via ArlH. However, ∼30 % of cells still form ArlD-GFP or ArlCE-GFP foci at the cell pole in the Δ*arlH* strains (Figures 5c, d, S4a, b), which indicates that ArlD and ArlCE can also position themselves at the cell pole by another mechanism. To test if the remainder of polar localization of ArlD and ArlCE, might be caused by interaction with other archaellum motor proteins, we created knock outs of ArlI and ArlJ in *H. volcanii*. ArlI is in direct interaction with ArlH, while ArlJ a membrane protein to which ArlI is bound (Banerjee *et al.*, 2013; Chaudhury *et al.*, 2016). For this reason, ArlH is not expected to bind to the archaellum motor in absence of either ArlI or ArlJ. Analysis on motility plate showed that the Δ*arlI* strain and the Δ*arl*J strain did not form motility rings (Fig S4C), which corresponds to the previously reported absence of archaella in similar knock-out strains in other archaea (Patenge *et al.*, 2001; Thomas *et al.*, 2002; Chaban *et al.*, 2007). Expression of GFP-ArlI in Δ*arlI* could restore the motile phenotype on semi-solid agar plates (Table S1) (Figure S4c). ArlI-GFP expression did not restore motility (Table S1). Expression of both fusion proteins was extremely low and could not be detected by Western blot analysis, while fluorescence microscopy showed a faint signal at the cell poles when GFP-ArlI was expressed in a Δ*arlI* background (Fig S4d).

Expression of ArlD-GFP in Δ*arlD* Δarl*I*, resulted in a reduction of the percentage of cells with polar foci (∼28%) compared to the expression of the same construct in the Δ*arlD* strain (∼70%) (Figure 5c). Similarly, when *arlJ* was deleted, 27% of cells kept polar ArlD-GFP foci. Thus, after deletion of either *arlH, arlI* and *arlJ* the number of cells with polar ArlD foci is strongly reduced but does not drop below 25%. Similar expression levels of ArlD-GFP were detected in all strains by Western blot analysis (Fig S1C). Comparable results were obtained for an ArlCE-GFP construct in a Δ*arlCE* Δ*arlH*, a Δ*arlCE* Δ*arlI* and a Δ*arlCE* Δ*arlJ* strain (Figure S4a, b). This suggests that positioning of ArlD and ArlCE at the cell poles, relies mainly on binding of these proteins to ArlH. However, a smaller population of ArlD and ArlCE proteins might also associate with other polar factors.

### Dynamics of ArlCE and ArlD

ArlD foci are dynamic in vivo and were previously shown to display mobility in the polar region of the cell (Li *et al.*, 2019) (Movie S1, Figure 6a). ArlD movement is mainly restricted to the cell pole, and occasionally also movement from particular small ArlD foci was observed from one cell pole to another (Li *et al.*, 2019). As our above described findings suggested that ArlCE and ArlD might form a precomplex required for interaction with the archaellum motor, we first made fluorescent time-lapse movies of ArlCE-GFP expressed in a *H. volcanii ΔarlCE* strain. Dynamic behavior of ArlCE was observed, specifically restricted to the cell pole. This movement of ArlCE was largely similar to the behavior of ArlD (Figure 6b, Movie S2). We assumed that this highly dynamic ArlD and ArlCE foci represent a part of the population that is not bound to the archaellum motor. To test this hypothesis, we studied the localization of the archaellum motor by time lapse microscopy of a Δ*arlH* strain expressing ArlH-GFP. ArlH foci remained localized at the cell pole over the course of an hour, suggesting that the archaellum motor is stably positioned at the cell pole (Movie S3, Figure 6c). This behavior is different from that of the dynamic behavior of ArlD and ArlCE. Together with the above described data, this suggests that ArlCE and ArlD likely form a complex, and that these complexes are localized at one or both cell poles. As the majority of the ArlCE and ArlD population is bound to the archaellum motor via ArlH, it is likely that specifically the unbound ArlCE and ArlD populations display the dynamics in the polar region.

**Figure 6.**
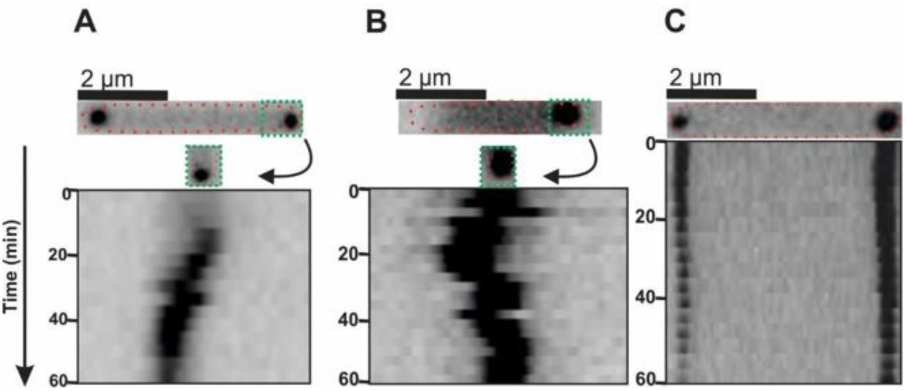
Intracellular dynamics of archaellum proteins ArlCE, ArlD and ArlH. Time-lapse images of (a) Δ*arlD* :: ArlD-GFP (b), Δ*arlCE ::* ArlCE-GFP (c) and Δ*arlH* GFP-ArlH. The upper panel shows a fluorescent image of the selected cells in which GFP-fused proteins were followed during 60 min. The lower panel displays kymographs of the cells shown on top. Cells are representable for >20 analyzed cells per strain. See also Movie S1, S2 and S3.

## Discussion

The motility structures of bacteria and archaea have a fundamentally different structural organization. Yet, both receive input from the chemotaxis system. This system allows cells to direct their movement along chemical gradients in order to find optimal conditions for survival. In bacteria, the central response regulator CheY binds to proteins at the base of the flagellum motor, the ‘switch complex’. As archaea lack homologs of the switch complex, we searched for archaellum motor proteins receiving signals from the chemotaxis system. Previously it has been shown that the archaeal specific adaptor protein CheF is required for functional chemotaxis in archaea. This protein can bind to archaeal CheY. However, the sequence order in which chemotactic signals are transferred via CheY and CheF to the archaeal motility machinery was until now unresolved. Our findings suggest that the archaellum proteins ArlCE and ArlD are the direct receivers of signals from the chemotaxis system and represent the archaeal equivalent of the ‘switch complex’.

By studying the localization of CheY-GFP fusions in different knock-out mutants, we could determine that in *H. volcanii*, CheY is primarily bound to the chemosensory arrays. This resembles the situation in *E.coli* where also the large majority of CheY proteins is present at the chemosensory arrays (Thiem *et al.*, 2007). In contrast, the polar positioning of CheF indicated that the largest fraction of this chemotaxis protein is normally bound to the archaellum motor.

Binding of CheF to the archaellum motor depends on the presence of both ArlCE and ArlD. Expression of fluorescent fusion proteins showed that ArlCE and ArlD co-localize and might form a complex. We suggest that the proteins form a precomplex, which is required for their binding to the archaellum motor. A strong interaction between these proteins would be in line with the fact that the *arlC, D* and *E* genes in different combinations are often fused in archaellum operons of different euryarchaea (Chaban *et al.*, 2007; Jarrell and Albers, 2012). In addition, a dependency of these proteins on each other would fit well with the hypothesizes that the ArlC, D and E proteins make up the unassigned density at the base of the archaellum motor in cryo-EM structures of euryarchaea (Daum *et al.*, 2017; Briegel *et al.*, 2017).

We found that the localization of both ArlCE and D depends on the archaellum motor protein ArlH (Figure 7). Also this is in line with the hypothesis that ArlCDE make up the ring like structure at the cytoplasmic side of the archaellum motor, since this part is in direct contact with the density in which the ArlH crystal structure could be mapped (Daum *et al.*, 2017).

**Figure 7.**
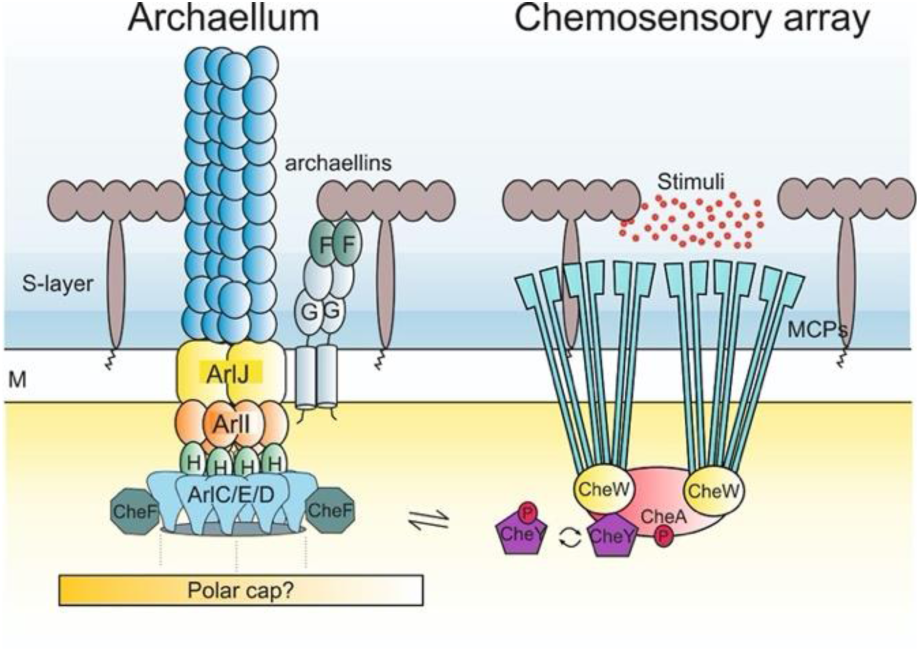
Simplified model of the chemotaxis signal transduction system and the archaellum in euryarchaea. MCPs are organized in chemosensory arrays together with CheW and CheA, which is necessary for signal integration and amplification. Autophosphorylation of CheA, results in phosphorylation of CheY. Phosphorylated CheY binds CheF, which is present at the base of the archaellum motor. CheF requires ArlCDE that form the switch complex, for binding to the archaellum motor. Possibly ArlCDE are not only binding to the archaellum motor protein ArlH, but are also interacting with the polar cap, to ensure polar localization. Chemotaxis accessory proteins are left out for simplicity. Blue, extracellular environment. Yellow, cytoplasm. M, membrane. MCP, methyl-accepting chemotaxis protein. Red dots, environmental stimuli that bind to the MCPs.

Surprisingly a small fraction of the ArlCE and ArlD proteins could still localize at the cell poles, even in absence of the archaellum motor (such as when ArlH, ArlI or ArlJ are deleted). Thus, ArlCE and ArlD are likely also capable of binding to other polar factors for their positioning. One candidate for this might be the ‘polar cap’ or ‘cytoplasmic cone’, which was revealed by whole-cell cryo-tomography of *P. furiosus* and *T. kodakaraensis* at a defined distance below the archaella motors and the cell membrane (Daum *et al.*, 2017; Briegel *et al.*, 2017). The polar cap is likely a typical feature of motile euryarchaeal cells (Kupper *et al.*, 1994; Daum *et al.*, 2017; Briegel *et al.*, 2017). The polar cap is in close proximity to the archaellum motor, but a direct connection was not observed (Daum *et al.*, 2017; Briegel *et al.*, 2017). The protein constitution of this polar cap is not known. It might be possible that ArlCE and ArlD can interact with the unknown polar cap protein(s) and as such are polar localized even in the absence of the archaellum. Precomplex formation might not be required for interaction with a pole organizing factor, as polar localization of GFP fused ArlD or ArlCE was still observed for a fraction of the cells, even when both *arlD* and *arlCE* were deleted. This is in contrast to binding of ArlD and ArlCE to the archaellum motor, which is strongly affected by the absence of either one of the proteins.

In summary, our findings suggest a central role for the ArlCE and ArlD proteins in the signal transduction from the chemotaxis system to the archaellum machinery. Extracellular stimuli are received by chemosensory receptors, organized in chemosensory arrays. Stimuli lead, via CheW, to autophosporylation of CheA and phosphorylation of CheY (Figure 7). Phosphorylated CheY then interacts with the archaeal specific chemotaxis protein CheF. CheF requires ArlCE and ArlD to bind to the archaellum motor. ArlCE and ArlD in their turn rely on interaction with the central motor protein ArlH (Figure 7). Possibly, a small proportion of the ArlCE and ArlD proteins also interacts with a polar organizing factor, such as the polar cap (Figure 7).

Thus, for a functional interaction between the bacterial-like chemotaxis system and the archaeal motility machinery, two important features are required in those archaea possessing both systems: i) the adaptor protein CheF and ii) ArlCDE that allow CheF binding to the archaellum motor. This work suggests that ArlCDE directly receive input from the chemotaxis system and might represent the archaeal equivalent of the switch complex. The hypothesis that they are conveniently located in the bell-like structure below the euryarchaeal archaellum motor, awaits further structural characterization of these three proteins.

## Experimental procedures

### Growth and genetic manipulation of *H. volcanii*

*H. volcanii* strains were cultured as previously described (Quax *et al.*, 2018b; Li *et al.*, 2019). Genetic manipulation and gene expression based on selection with uracil in Δ*pyrE2* strains were performed with PEG 600 as described previously (Allers *et al.*, 2004). The cells were cultured shaking at 120 rpm at 45 °C or 42 °C in complete YPC medium containing 5% Bacto™yeast extract, 1% peptone (Oxoid, UK), 1% BactoTM Casamino acids (BD Biosciences, UK) or in selective CA medium containing only 5% BactoTM casamino acids in 18 % SW (Salt water, containing per liter 144 g NaCl, 21 g MgSO_4_ × 7H_2_O, 18 g MgCl_2_ × 6H_2_O, 4.2 g KCl, and 12 mM Tris HCl, pH 7.3). Plasmids based on pTA1228 (Brendel *et al.*, 2014), with *pyrE2* for selection with uracil, were constructed to express proteins in *H. volcanii* strains (Table S3). Plasmids based on pTA131 were used to create knock-out constructs for the pop-in pop-out method based on the pyrE2 gene. Salt stable GFP and mCherry genes were introduced to pTA1228 plasmid, allowing expression of N-terminal and C-terminal fluorescent fusion proteins in *H. volcanii* strains (Duggin et al, Nature, 2015).

### Deletions of genes in *H. volcanii* strains

The primers used to create knockout plasmids based on pTA131 are described in Table S2. Construction and transformation of knock-out plasmids was performed as described previously (Allers *et al.*, 2004). Selection for pop-in occurred on CA plates. This was followed by 3 transfers in liquid YPC medium. Then the inoculum was diluted 10^−3^ and cultured on CA plates with 50 μg/mL 5-FOA and 10 μg/mL uracil for the pop-out selection. Colonies were streaked on a new YPC plate, grown for two days, and subjected to a colony lift to Zeta-Probe℗ GT Blotting membranes (Biorad). After cell lysis and DNA cross-linking, the DNA was subjected to pre-hybridization and hybridization using a DIG High Prime DNA labeling and detection starter kit II (Roche) according to the manufacturer’s instructions with a DIG-labeled probe of ∼100-200 bp annealing in the targeted gene (for primer sequences, see Table S2). Colonies to which the probe did not bind were grown in liquid YPC media, and genomic DNA was isolated as described previously (Allers *et al.*, 2004). The genomic DNA of several selected mutants was analyzed with PCR using primers that anneal outside of the flanking regions of the deleted gene (see Table S2), and the products formed were compared with those of the wild-type H26 strain on an agarose gel.

### Strains, plasmids and primers

The strains, plasmid and primer sequences used in this study can be found in Tables S2-S4.

### Motility assays of *H. volcanii* on semisolid agar plates

Motility assays were performed using the same method as previously described (Quax *et al.*, 2018b; Li *et al.*, 2019). Semi solid agar plates were made in YPC medium containing 0.3% agar, 50 μg/mL uracil and 1 mM tryptophan. Fresh cells were inoculated in 5 mL CA medium with 50 μg/mL uracil and/or tryptophan when required. 10µL of the inoculum of each strain were dropped on the same semi solid agar plates. The experiment was performed at least three independent times (containing 3 biological replicates each). The motility ring formed on semi solid agar plates after cultivation at 45 °C for 5 days were scanned, and the diameter was measured.

### Transmission Electron Microscopy

Cells were grown over night at 42 °C in CA medium to an OD600 of ∼0.05 and then harvested by centrifugation of 3000 g for 10 min. Cells were concentrated and resuspended in CA medium with 2% (vol/vol) glutaraldehyde and 1% (vol/vol) para-formaldehyde. Cells were adsorbed to glow-discharged carbon-coated grids with Formvar films for 30 sec. Samples were washed three times in distilled H2O and negatively stained with 2% (wt/vol) uranyl acetate. Cells were imaged using a Philips CM10 transmission EM coupled to a Gatan BioScan camera and the Gatan DigitalMicrograph software.

### Western blotting analysis

Strains were grown under similar conditions as those for fluorescence microscopy analysis. The cells were harvested by centrifugation and concentrated to a theoretical OD of 22. Total cell lysates were separated on by SDS-PAGE (sodium dodecyl sulfate–polyacrylamide gel electrophoresis) on two 12% acrylamide gels. One gel was blotted to a PVDF (Polyvinylidene fluoride) membrane (Roche) by semi-dry Western blotting. The gels were stained with Quick Coomassie (Generon Ltd.). The membrane was subsequently incubated in blocking buffer (0.2% I-Block™, 0.1% Tween) (Thermo Fisher Scientific, MA, USA) at room temperature for 2 h. After 3 times washing with PBST buffer (0.1 g/L CaCl_2_, 0.2 g/L KCl, 0.2 g/L KH2PO4, 0.1 g/L MgCl_2_×6H_2_O, 8 g/L NaCl, 1.15 g/L Na_2_HPO_4_, pH 7.4), the membrane was incubated in anti-GFP antibody diluted to 1:1,000 in PBST buffer for 1 h at room temperature. After three washes in PBST, a secondary anti-rabbit antibody (from goat) coupled to HRP (horseradish peroxidase) (Thermo Fisher Scientific, MA, USA) was added to the membrane (1:5,000). Clarity ECL Western blotting substrate (Thermo Fisher Scientific) was used for visualization of the chemiluminescent signals in a ECL ChemoCam Imager (Intas, Germany).

### Fluorescence Microscopy

*H. volcanii* precultures were grown in 5 mL CA medium overnight. The next day the cultures were diluted to a theoretical OD of 0.005 in 20 mL of CA medium at 42 °C. After 16 hours the cultures had an OD of 0.01-0.05 and were imaged. During the last hour before observation by microscopy, 0.2 mM tryptophan was added. Cells were pipetted on an agarose pad (1% agar, 18% SW) and covered with a glass slip. The cells were observed at 100x magnification in the phase contrast (PH3) mode on a Zeiss Axio Observer 2.1 Microscope equipped with a heated XL-5 2000 Incubator running VisiVIEW℗ software. Each experiment was repeated on at least three independent occasions resulting in the analysis of over 500 cells per strain.

To track the mobility of protein foci with live imaging, 0.38% agar pads made of CA containing 1 mM tryptophan were poured in a round DF 0.17 mm microscopy dish (Bioptechs). After drying, the cells were placed underneath the agar pad, and the lid was placed on the microscopy dish. Images in the PH3 and GFP modes were captured at 100 x magnification every 3 minutes for 1 hour at 45 °C.

### Image analysis

Microscopy images were processed using the ImageJ plugin MicrobeJ (Ducret *et al.*, 2016). The number of foci per cell was counted, and the cells were divided over bins based on the number of cellular foci. The number of cells with the same positioning patterns was calculated as a percentage of the total. The parameters of the detection of the fluorescent foci were set to the same levels for all the proteins analyzed. Fluorescent foci movement in the time-lapse image series was characterized by time-space plots generated by the ‘Surface plotter’ function in ImageJ (Ducret *et al.*, 2016). To determine if the percentage of cells with foci was significantly different between strains, an unpaired two-tailed T-test was performed on the percentages calculated for each independent experiment (minimally 3). Total number of analyzed cells was >500 per strain.

## Acknowledgements

We thank Frank Braun and Phillip Nußbaum for support with experiments. The TEM is operated by the University of Freiburg, Faculty of Biology, as a partner unit within the Microscopy and Image Analysis Platform, Freiburg. This work was supported by the Deutsche Forschungsgemeinschaft (German Research Foundation) with an Emmy Nöther grant (411069969) to T.E.F.Q. and a grant within the Collaborative Research Centre SFB 1381 (403222702-SFB 1381) S-.V.A. Z.L. was supported by a CSC scholarship from the Chinese government.

## Authors Contribution

T.E.F.Q. and S.-V.A. designed research. Z.L., M.R. and T.E.F.Q. performed research and interpreted data. T.E.F.Q. and S.-V.A. wrote the paper. All authors read and contributed to the manuscript.

## Data availability statement

The data that supports the findings of this study are available in the supplementary material of this article

## Abbreviated Summary

The bacterial-like chemotaxis system is essential for directional movement in Archaea. So far, it was unknown how the signal is transferred from the archaeal CheY to the archaellum motor to initiate motor switching. In this study we demonstrate that the proteins ArlCDE represent the archaellum switch complex which is the docking point for the CheY-CheF complex and essential for archaellum filament switching.

