## Supplemental Material for "ArlCDE form the archaeal switch complex"

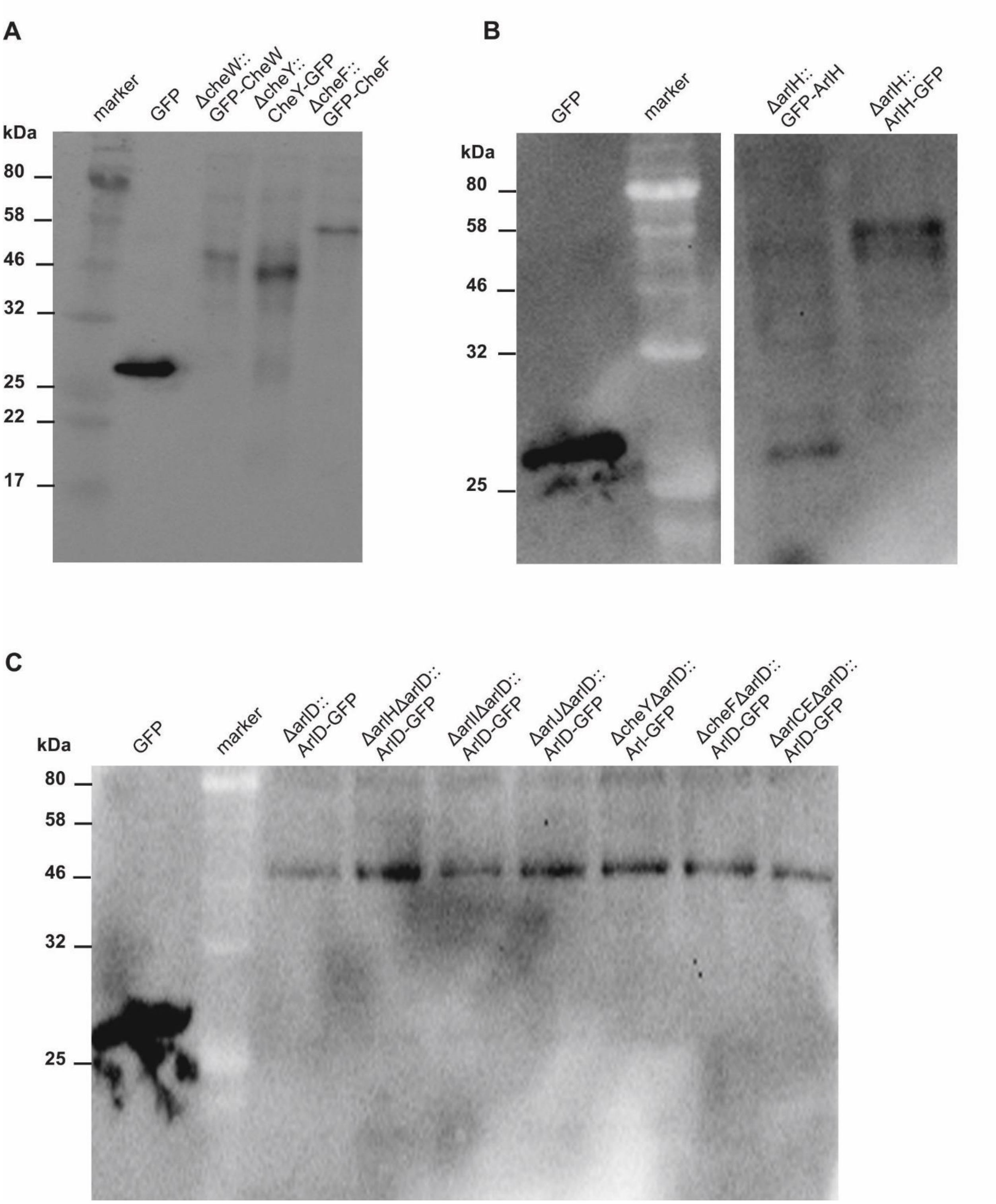


**Figure S1. α-GFP Western blot** on total cell lysates of strains expressing GFP-fused constructs. (a) Western blot of single knock-out strains expressing GFP fusions with CheW, CheY and CheF. (b) Western blot of a Δ*arlH* strain expressing N and C terminal fusions to ArlH. For GFP-ArlH the GFP monomer was mainly detected, indicating that there is likely cleavage between GFP and ArlH (c) ArlD-GFP expression in multiple double deletion strains, showing that the fusion protein is correctly expressed in all backgrounds. Arrows indicate detected GFP fusions. Marker, protein marker with known sizes. GFP, positive control of purified GFP protein (27 kDa). GFP-CheW (44.3 kDa), GFP-CheF (60 kDa), CheY-GFP (42 kDa), ArlH-GFP (55 KDa), ArlD-GFP (48.5 kDa).

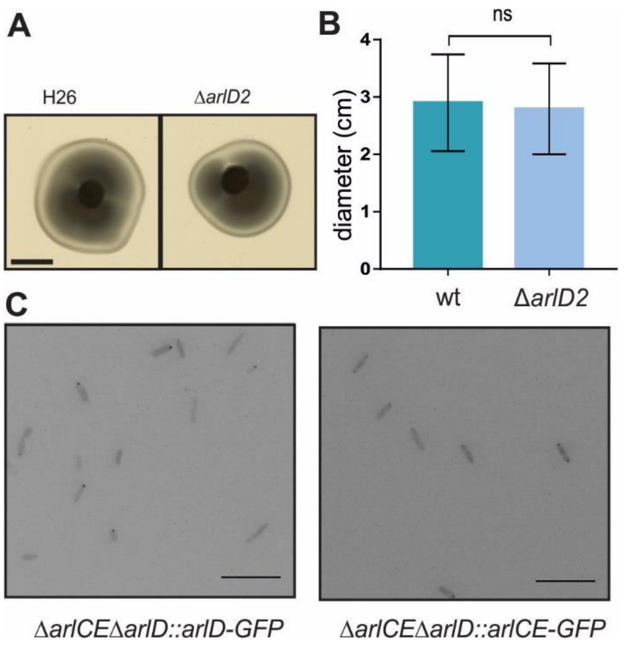


**Figure S2. Positioning patterns of ArlD and ArlCE are dependent on each other.** (a) Representative example of analysis of an Δ*arlD2 H. volcanii* strain on semi-solid agar plate in comparison with a control strain H26, showing that ArlD2 does not affect directional movement. (b) Quantification of diameters of motility rings on semi-solid agar plates, such as shown in panel (a). Experiment is performed on at least 3 independent occasions with 3 biological replicates each. Ns, not significant (c) Left: Intracellular distribution of ArlD-GFP in a *H. volcanii* Δ*arlCE* Δ*arlD* strain. Right: Intracellular distribution of ArlCE-GFP foci in *H. volcanii* Δ*arlCE* Δ*arlD* strain. Scale bars, 10µm.

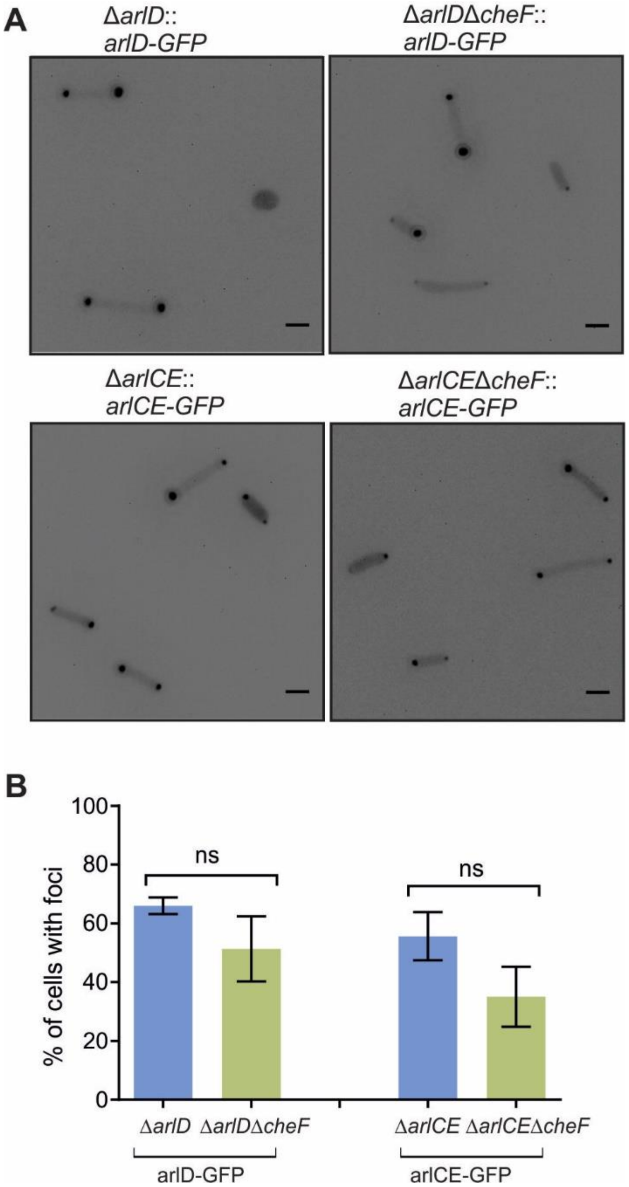


**Figure S3. CheF is not required for intracellular positioning of ArlD and ArlCE.** (a) Representative fluorescent images of intracellular distribution of ArlD and ArlCE in the absence of CheF. Scale bars, 2 µm. (b) Percentages of cells with intracellular foci in the strains described in (a). n>500. ns, not significant P>0.01 as determined by T-test.

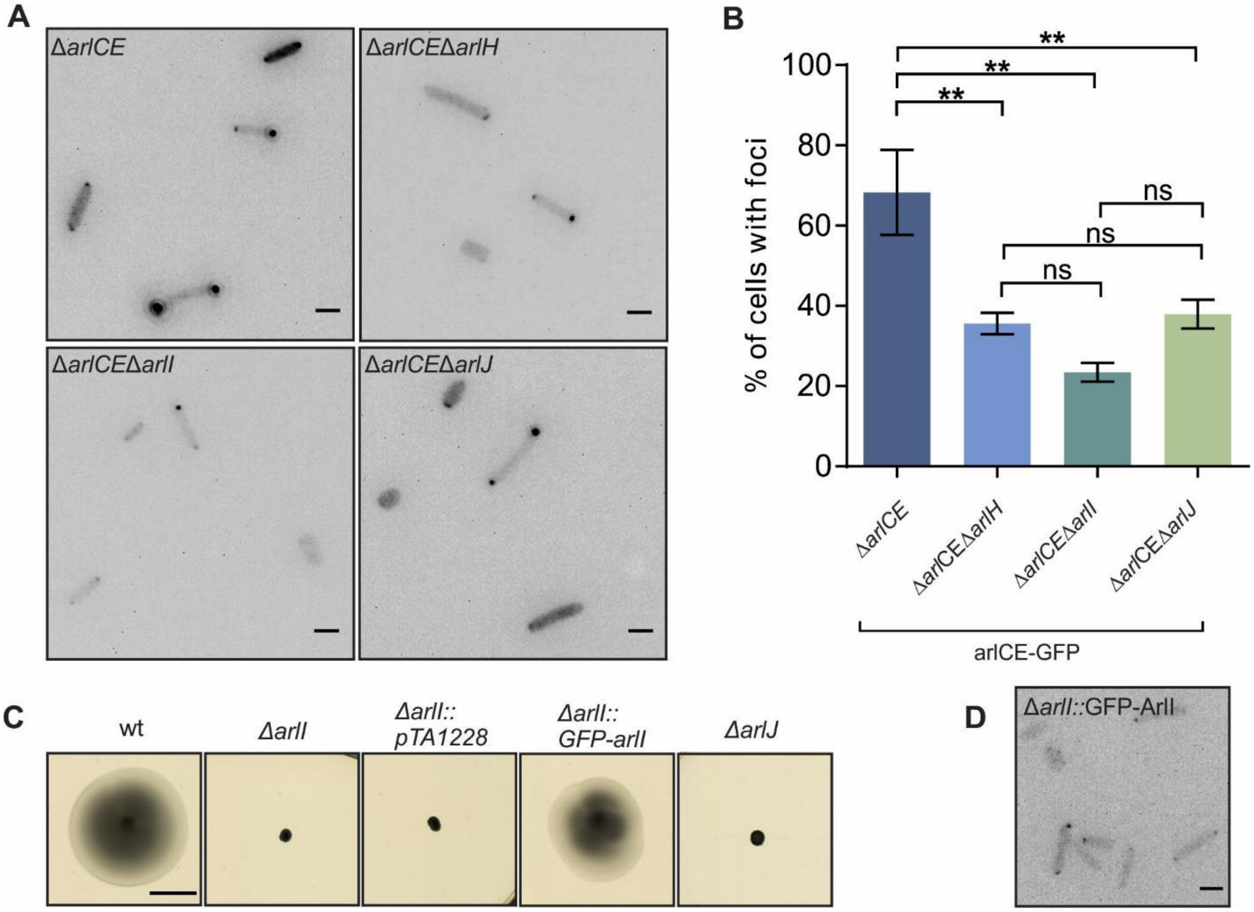


**Figure S4. Intracellular distribution of ArlCE in the absence of motor proteins in *H. volcanii.*** (a) Intracellular distribution of ArlCE-GFP clusters in *H. volcanii* strains in which different archaellum motor genes were deleted. Scale bars, 2 µm. (b) Percentages of cells with intracellular ArlCE-GFP foci in several strains. **, $P < 0.01$, ns, not significant, $P>0.05$. (c) Influence of ArlI and ArlJ on directional movement as assayed by semi-solid agar plate. Scale bar, 1 cm. (d) Representative fluorescent image of GFP-ArlI expression in a Δ*arlI* background, showing faint polar foci. Scale bar, 2 µm.

### Supplemental Tables

**Table S1** Analysis of localization pattern of several GFP-fused motility and chemotaxis proteins in *H. volcanii*

| Mutants | Restoration of motile phenotype of mutant by expressed protein | | | Localization pattern fusion protein | | References |
|---|---|---|---|---|---|---|
| | Untagged | C-term | N-term | C-term | N-term | |
| ***ΔarlI*** | Yes | Yes | No | 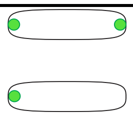 | 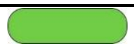 | This study |
| ***ΔarlH*** | Yes | Yes* | No | 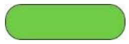 | 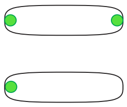 | This study |
| ***ΔarlD*** | Yes | No | Yes | 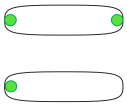 | 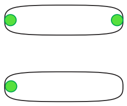 | Li et al, 2019 |
| ***ΔarlCE*** | Yes | No | No | 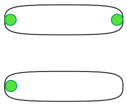 | 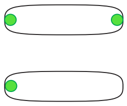 | This study |
| ***ΔcheF*** | Yes | Yes | No | 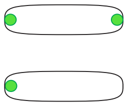 | 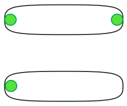 | Li et al, 2019 |
| ***ΔcheY*** | Yes | No | Yes | 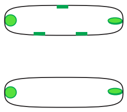 | 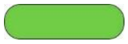 | Li et al, 2019 |
| ***ΔcheW*** | Yes | Yes | No | 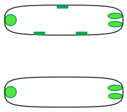 | 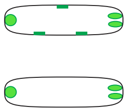 | Li et al, 2019 |

*Western blot showed that the fusion protein was not correctly expressed

### Table S2 – Primers used in this study

| Name | Sequence | Description |
|---|---|---|
| **9028** | GGGGTACCACCCTCATCGAACTGGTCG | forward primer with KpnI site for amplification of upstream flank of *arlCE* of *H. volcanii* for knock-out plasmid pTA131 |
| **9029** | GGAATTCCATATGTTGTTCTTCCGTGTCCG TTTC | reverse primer with NdeI site for amplification of upstream flank of *arlCE* of *H. volcanii* for knock-out plasmid pTA131 |
| **9030** | GGAATTCCATATGGGATTCGGTGTTAGC | forward primer with NdeI site for amplification of downstream flank of *arlCE* of *H. volcanii* for knock-out plasmid pTA131 |
| **9031** | CTAGTCTAGAGTCACGGTATCGATGAG | reverse primer with XbaI site for amplification of downstream flank of *arlCE* of *H. volcanii* for knock-out plasmid pTA131 |
| **9057** | CGCCATATGTCATCAGACGATGGACTTGGA TTC | forward primer for amplification of *arlCE* from *H.volcanii* with NdeI site for cloning in pIDJl-40 |
| **9004** | CGGGATCCACACCGAATCCCATCGAATCT CC | reverse primer for amplification of *arlCE* from *H.volcanii* with BamHI site for cloning in pIDJl-40 |
| **9058** | CTAGCTAGCATGTCATCAGACGATGGACTT G | forward primer for amplification of *arlCE* from *H.volcanii* with NheI site for cloning in pSVA3922 |
| **7188** | CGGGATCCCTAACACCGAATCCCATCGAA TCTCC | reverse primer for amplification of *arlCE* from *H.volcanii* with BamHI site for cloning in pSVA3922 |
| **9003** | GGAATTCCATATGGTGCTCGCCGCGAGCA TGATG | forward primer for amplification of *arlCE* from *H.volcanii* with NdeI site for cloning in pTA1392 |
| **9023** | CGGGATCCCTAACACCGAATCCCATCGAA TCTCC | reverse primer for amplification of *arlCE* from *H.volcanii* with BamHI site for cloning in pTA1392 |
| **7128** | GGAATTCCATATGTACCTGGACCCGGACG AATACGACCCAG | forward primer for amplification of *arlD1* from *H.volcanii* with NdeI site for cloning in pSVA3943 |
| **7131** | CTAGCTAGCCACCATCGAGGAGAGGCGGG C | reverse primer for amplification of *arlD1* from *H.volcanii* with NheI site for cloning in pSVA3943 |

| | | |
|---|---|---|
| **7221** | ATTGGATCCGTGCTCGCCGCGAGCATGAT | forward primer for amplification of *arlCE* from *H.volcanii* with BamHI site for cloning in pSVA5072 |
| **7222** | CGGATGCATACACCGAATCCCATCGAATCTCCCC | reverse primer for amplification of *arlCE* from *H.volcanii* with NsiI site for cloning in pSVA5072 |
| **7262** | GGGGTACCTCTCGTTCGTCGTGGACGCC | forward primer with KpnI site for amplification of upstream flank of *arlH* of *H. volcanii* for knock-out plasmid pTA131 |
| **10571** | GTCTCCGACCTGCTGGCCCATCATGGTACGCGGAATTCGAACACCTC | reverse primer for amplification of upstream flank of *arlH* of *H. volcanii* for knock-out plasmid pTA131 |
| **10572** | TGGGCCAGCAGGTCGGAGACCGCGTCG | forward primer for amplification of downstream flank of *arlH* of *H. volcanii* for knock-out plasmid pTA131 |
| **7265** | GCTCTAGAGGCCGACGATGTCGCGGACGAGTCG | reverse primer with XbaI site for amplification of downstream flank of *arlH* of *H. volcanii* for knock-out plasmid pTA131 |
| **7216** | GGAATTCCATATGAGCAGTCAATTCTCTCTCGGGCTATCCG | forward primer for amplification of *arlH* from *H. volcanii* introducing NdeI site for cloning in pTA1392 |
| **7215** | GGAATTCCTACGCCACGCTCCTCGACTC | reverse primer for amplification of *arlH* from *H. volcanii* introducing EcoRI site and stop codon for cloning in pTA1392 |
| **9027** | GGAATTCCATATGATGGAGTGGCAGAACGACGAAG | forward primer for amplification of *arlH* from *H.volcanii* with BamHI site for cloning in pTA1392 |
| **9083** | CGGGGTACCGACATCGTCAAGCATCTTCTC | forward primer with KpnI site for amplification of upstream flank of *arlI* of *H. volcanii* for knock-out plasmid pTA131 |
| **9084** | CCGGAATTCCTACGCCACGCTCCTCGAC | reverse primer for amplification of upstream flank of *arlI* of *H. volcanii* for knock-out plasmid pTA131 |
| **9085** | CCGGAATTCCGAGATGGCAGGGGAACAGAG | forward primer for amplification of downstream flank of *arlI* of *H. volcanii* for knock-out plasmid pTA131 |

| | | |
|---|---|---|
| **9086** | TGCTCTAGAAGGAAGTCGTCGATGCTCTG | reverse primer with XbaI site for amplification of downstream flank of *arlI* of *H. volcanii* for knock-out plasmid pTA131 |
| **10561** | CGGGGTACCGCGTCCGACATCGTCTCGATGATTC | forward primer with KpnI site for amplification of upstream flank of *arlJ* of *H. volcanii* for knock-out plasmid pTA131 |
| **10562** | TCTGTGGTCCGTCGTCAGTCATCTCGTTACACCCTCGCCATGTCGAACGG | reverse primer for amplification of upstream flank of *arlJ* of *H. volcanii* for knock-out plasmid pTA131 |
| **10563** | ATGACTGACGACGGACCACAGAG | forward primer for amplification of downstream flank of *arlJ* of *H. volcanii* for knock-out plasmid pTA131 |
| **10564** | TGCTCTAGAGCGGAACAACAGCAGCGAGAAC | reverse primer with XbaI site for amplification of downstream flank of *arlJ* of *H. volcanii* for knock-out plasmid pTA131 |
| **10552** | CGGAATTCATGGCGACCGAACACGGCTC | forward primer for amplification of *arlI* from *H.volcanii* with EcoRI site for cloning in pIDJl-40 |
| **10553** | CGCGGATCCCACCCTCGCCATGTCGAACGG | reverse primer for amplification of *arlI* from *H.volcanii* with BamHI site for cloning in pIDJl-40 |
| **10554** | CTAGCTAGCATGGCGACCGAACACGGCTC | forward primer for amplification of *arlI* from *H.volcanii* with NheI site for cloning in pSVA3922 |
| **10555** | CGCGGATCCTTACACCCTCGCCATGTCGAACGG | reverse primer for amplification of *arlI* from *H.volcanii* with BamHI site for cloning in pSVA3922 |

**Table S3 – Plasmids used in this study**

| Name | Description | Source |
|---|---|---|
| **pTA131** | pyrE2 marked deletion plasmid *H.volcanii*. Amp resistance. | (Allers *et al.*, 2004) |
| **pTA1228** | pyrE2 marked protein expression plasmid *H.volcanii*. Amp resistance. Tryptophan inducible. | (Brendel *et al.*, 2014) |
| **pIDJL-40** | Expression plasmid *H.volcanii* for C-terminal GFP tagging based on pTA1228. | (Duggin *et al.*, 2015) |
| **pSVA3922** | Expression plasmid *H.volcanii* for N-terminal GFP tagging based on pTA1228. | (Li *et al.*, 2019) |
| **pSVA5058** | pTA131 with upstream and downstream flanks of cheF1. Deletion plasmid to delete the cheF1 gene in *H.volcanii*. Deletion cassette is made by ligation via BamHI site of ~500 bp upstream and downstream flanking region. | (Quax *et al.*, 2018) |
| **pSVA5611** | pIDJL-40 with *H. volcanii* cheY. Expression plasmid to express CheY-GFP under trp promoter. | (Li *et al.*, 2019) |
| **pSVA5029** | pTA131 with upstream and downstream flanks of cheW1. Deletion plasmid to delete the cheW1 gene in *H.volcanii*. Deletion cassette is made by ligation via BamHI site of ~500 bp upstream and downstream flanking region. | (Li *et al.*, 2019) |
| **pSVA5079** | pSVA3922 with *H. volcanii* cheF1. Expression plasmid to express GFP-CheF1 under trp promoter. | (Li *et al.*, 2019) |
| **pSVA5004** | pTA131 with upstream and downstream flanks of ArlD1. Deletion plasmid to delete the ArlD1 gene in *H.volcanii*. Deletion cassette is made by ligation via BamHI site of ~500 bp upstream and downstream flanking region. | (Li *et al.*, 2019) |
| **pSVA5610** | pTA131 with upstream and downstream flanks of ArlCE. Deletion plasmid to delete the ArlCE gene in | This study |

| | | |
|---|---|---|
| | *H.volcanii*. Deletion cassette is made by ligation via BamHI site of ~500 bp upstream and downstream flanking region. | |
| **pSVA5604** | pIDJL-40 with *H. volcanii* ArlCE. Expression plasmid to express ArlCE-GFP under trp promoter. | This study |
| **pSVA5626** | pSVA3922 with *H. volcanii* ArlCE. Expression plasmid to express GFP-ArlCE under trp promoter. | This study |
| **pSVA5606** | pSVA1228 with *H. volcanii* ArlCE. Expression plasmid to express untagged ArlCE under trp promoter. | This study |
| **pSVA3943** | Double expression plasmid dimeric GFP-mCherry both C-terminal fusions based on pTA1228. | This study |
| **pSVA5072** | pSVA3943 with *H. volcanii* ArlD1. Cloning intermediate to create pSVA5073. | This study |
| **pSVA5073** | pSVA5072 with *H. volcanii* ArlCE. Expression plasmid expression plasmid to co-express ArlD1-GFP and ArlCE-mCherry under trp promoter. | This study |
| **pSVA3919** | pIDJL-40 with *H. volcanii* ArlD1. Expression plasmid to express ArlD1-GFP under trp promoter. | (Li *et al.*, 2019) |
| **pSVA5669** | pTA131 with upstream and downstream flanks of ArlH. Deletion plasmid to delete the ArlH gene in *H.volcanii*. Deletion cassette is made by fusion of of ~500 bp upstream and downstream flanking region via overlap PCR. | This study |
| **pSVA5608** | pSVA1228 with *H. volcanii* ArlH. Expression plasmid to express untagged ArlH under trp promoter. | This study |
| **pSVA5068** | pSVA3922 with *H. volcanii* ArlH. Expression plasmid to express GFP-ArlH under trp promoter. | This study |
| **pSVA5609** | pIDJL-40 with *H. volcanii* ArlH. Expression plasmid to express ArlH-GFP under trp promoter. | This study |
| **pSVA5637** | pTA131 with upstream and downstream flanks of ArlI. Deletion plasmid to delete the ArlI gene in *H.volcanii*. | This study |

| | | |
|---|---|---|
| | Deletion cassette is made by fusion of of ~500 bp upstream and downstream flanking region via overlap PCR. | |
| **pSVA5667** | pTA131 with upstream and downstream flanks of ArlJ. Deletion plasmid to delete the ArlJ gene in *H.volcanii*. Deletion cassette is made by fusion of of ~500 bp upstream and downstream flanking region via overlap PCR. | This study |
| **pSVA5662** | pIDJL-40 with *H. volcanii* ArlI. Expression plasmid to express ArlI-GFP under trp promoter. | This study |
| **pSVA5663** | pSVA3922 with *H. volcanii* ArlI. Expression plasmid to express GFP-ArlI under trp promoter. | This study |

### Table S4 – Strains used in this study

#### Strains

| Name | Organism | Background strain | Genotype | Used plasmid | Source |
|---|---|---|---|---|---|
| **H26** | *H.volcanii* | WT | Δ*pyrE2* | | (Allers *et al.*, 2004) |
| **HTQ32** | *H.volcanii* | H26 | Δ*pyrE2* Δ*cheY* | | (Quax *et al.*, 2018) |
| **HTQ392** | *H.volcanii* | HTQ32 | Δ*pyrE2* Δ*cheF1* Δ*cheY* | pSVA5058 | This study |
| **HTQ581** | *H.volcanii* | HTQ392 | Δ*pyrE2* Δ*cheF1* Δ*cheY* Δ*cheW1* | pSVA5029 | This study |
| **HTQ364** | *H.volcanii* | HTQ32 | Δ*pyrE2* Δ*cheY ::[cheY -GFP]* | pSVA5611 | (Li *et al.*, 2019) |
| **HTQ559** | *H.volcanii* | HTQ392 | Δ*pyrE2* Δ*cheF1* Δ*cheY::[cheY -GFP]* | pSVA5611 | This study |
| **HTQ590** | *H.volcanii* | HTQ581 | Δ*pyrE2* Δ*cheF1* Δ*cheY* Δ*cheW1* *::[cheY -GFP]* | pSVA5611 | This study |
| **HTQ403** | *H.volcanii* | H26 | Δ*pyrE2* Δ*cheF1* | | (Quax *et al.*, 2018) |
| **HTQ355** | *H.volcanii* | HTQ403 | Δ*pyrE2* Δ*cheF1::[GFP-cheF1]* | pSVA5079 | This study |
| **HTQ558** | *H.volcanii* | HTQ392 | Δ*pyrE2* Δ*cheF1* Δ*cheY::[GFP-cheF1]* | pSVA5079 | This study |
| **HTQ384** | *H.volcanii* | HTQ403 | Δ*pyrE2* Δ*arlD1* Δ*cheF1* | pSVA5004 | This study |
| **HTQ390** | *H.volcanii* | HTQ403 | Δ*pyrE2* Δ*arlCE* Δ*cheF1* | pSVA5610 | This study |
| **HTQ331** | *H.volcanii* | HTQ384 | Δ*pyrE2* Δ*arlD1* Δ*cheF1::[GFP-cheF1]* | pSVA5079 | This study |
| **HTQ371** | *H.volcanii* | HTQ390 | Δ*pyrE2* Δ*arlCE* Δ*cheF1::[GFP-cheF1]* | pSVA5079 | This study |
| **HTQ360** | *H.volcanii* | H26 | Δ*pyrE2* Δ*ArlCE* | pSVA5610 | This study |
| **HTQ369** | *H.volcanii* | HTQ360 | Δ*pyrE2* Δ*arlCE::[GFP-arlCE]* | pSVA5626 | This study |

| | | | | | |
|---|---|---|---|---|---|
| **HTQ370** | *H.volcanii* | HTQ360 | Δ*pyrE2* Δ*arlCE::[ arlCE-GFP]* | pSVA5604 | This study |
| **HTQ376** | *H.volcanii* | HTQ360 | Δ*pyrE2* Δ*arlCE::[PyrE2+]* | pTA1228 | This study |
| **HTQ377** | *H.volcanii* | HTQ360 | Δ*pyrE2* Δ*arlCE::[ arlCE]* | pSVA5606 | This study |
| **HTQ19** | *H.volcanii* | H26 | Δ*pyrE2* Δ*arlD1* | | (Li *et al.*, 2019) |
| **HTQ207** | *H.volcanii* | HTQ19 | Δ*pyrE2* Δ*arlD1::[arlD1-GFP]* | pSVA3919 | (Li *et al.*, 2019) |
| **HTQ380** | *H.volcanii* | HTQ19 | Δ*pyrE2* Δ*arlD1* Δ*arlCE* | pSVA5610 | This study |
| **HTQ307** | *H.volcanii* | HTQ380 | Δ*pyrE2* Δ*arlD1* Δ*arlCE::[arlD1-GFP _arlCE-mCherry]* | pSVA5073 | This study |
| **HTQ308** | *H.volcanii* | HTQ380 | Δ*pyrE2* Δ*ArlD1* Δ*arlCE::[arlCE-GFP]* | pSVA5604 | This study |
| **HTQ309** | *H.volcanii* | HTQ380 | Δ*pyrE2* Δ*arlD1* Δ*arlCE::[arlD1-GFP]* | pSVA3919 | This study |
| **HTQ330** | *H.volcanii* | HTQ384 | Δ*pyrE2* Δ*arlD1* Δ*cheF1::[arlD1-GFP]* | pSVA3919 | This study |
| **HTQ372** | *H.volcanii* | HTQ390 | Δ*pyrE2* Δ*arlCE* Δ*cheF1::[arlCE-GFP]* | pSVA5604 | This study |
| **HTQ570** | *H.volcanii* | H26 | Δ*pyrE2* Δ*arlH* | pSVA5669 | This study |
| **HTQ396** | *H.volcanii* | H26 | Δ*pyrE2* Δ*arlI* | pSVA5637 | This study |
| **HTQ574** | *H.volcanii* | H26 | Δ*pyrE2* Δ*arlJ* | pSVA5667 | This study |
| **HTQ572** | *H.volcanii* | HTQ570 | Δ*pyrE2* Δ*arlH::[arlH]* | pSVA5608 | This study |
| **HTQ579** | *H.volcanii* | HTQ570 | Δ*pyrE2* Δ*arlH::[GFP-arlH]* | pSVA5068 | This study |
| **HTQ580** | *H.volcanii* | HTQ570 | Δ*pyrE2* Δ*arlH::[arlH-GFP]* | pSVA5069 | This study |
| **HTQ556** | *H.volcanii* | HTQ396 | Δ*pyrE2* Δ*ArlI::[ArlI-GFP]* | pSVA5662 | This study |
| **HTQ557** | *H.volcanii* | HTQ396 | Δ*pyrE2* Δ*arlI ::[GFP-arlI]* | pSVA5663 | This study |
| **HTQ397** | *H.volcanii* | HTQ19 | Δ*pyrE2* Δ*arlD1* Δ*arlI* | pSVA5637 | This study |
| **HTQ398** | *H.volcanii* | HTQ360 | Δ*pyrE2* Δ*arlCE* Δ*arlI* | pSVA5637 | This study |
| **HTQ575** | *H.volcanii* | HTQ19 | Δ*pyrE2* Δ*arlD1* Δ*arlJ* | pSVA5667 | This study |
| **HTQ576** | *H.volcanii* | HTQ360 | Δ*pyrE2* Δ*arlCE* Δ*arlJ* | pSVA5667 | This study |
| **HTQ584** | *H.volcanii* | HTQ19 | Δ*pyrE2* Δ*arlD1* Δ*arlH* | pSVA5669 | This study |

| | | | | | |
|---|---|---|---|---|---|
| **HTQ585** | *H.volcanii* | HTQ360 | Δ*pyrE2 ΔarlCE ΔarlH* | pSVA5669 | This study |
| **HTQ554** | *H.volcanii* | HTQ397 | Δ*pyrE2 ΔarlD1 ΔarlI:: [arlD1-GFP]* | pSVA3919 | This study |
| **HTQ555** | *H.volcanii* | HTQ398 | Δ*pyrE2 ΔarlCE ΔarlI :: [arlCE-GFP]* | pSVA5604 | This study |
| **HTQ582** | *H.volcanii* | HTQ575 | Δ*pyrE2 ΔarlD1 ΔarlJ:: [arlD1-GFP]* | pSVA3919 | This study |
| **HTQ583** | *H.volcanii* | HTQ576 | Δ*pyrE2 ΔarlCE ΔarlJ :: [arlCE-GFP]* | pSVA5604 | This study |
| **HTQ393** | *H.volcanii* | HTQ384 | Δ*pyrE2 ΔarlD1 ΔarlH:: [arlD1-GFP]* | pSVA3919 | This study |
| **HTQ394** | *H.volcanii* | HTQ385 | Δ*pyrE2 ΔarlCE ΔarlH :: [arlCE-GFP]* | pSVA5604 | This study |

**Supplemental Movies**

Movie S1: Time lapse movie of 1 h of an Δ*arlD* strain expressing ArlD-GFP in *H. volcanii*

Movie S2: Time lapse movie of 1 h of an Δ*arlCE* strain expressing ArlCE-GFP in *H. volcanii*

Movie S3: Time lapse movie of 1 h of an Δ*arlH* strain expressing ArlH-GFP in *H. volcanii*
